# Guilty by association: How group-based (collective) guilt arises in the brain

**DOI:** 10.1101/683730

**Authors:** Zhiai Li, Hongbo Yu, Yongdi Zhou, Tobias Kalenscher, Xiaolin Zhou

## Abstract

People do not only feel guilty for transgressions of social norms/expectations that they are causally responsible for, but they also feel guilty for transgressions committed by those they identify as in-group (i.e., collective or group-based guilt). However, the neurocognitive basis of group-based guilt and its relation to personal guilt are unknown. To address these questions, we combined functional MRI with an interaction-based minimal group paradigm in which participants either directly caused harm to victims (i.e., personal guilt), or observed in-group members cause harm to the victims (i.e., group-based guilt). In three experiments (N = 90), we demonstrated that perceived shared responsibility with in-group members in the transgression predicted behavioral and neural manifestations of group-based guilt. Multivariate pattern analysis of the functional MRI data showed that group-based guilt recruited a similar brain representation in anterior middle cingulate cortex as personal guilt. These results have broaden our understanding of how group membership is integrated into social emotions.

## Introduction

Guilt is viewed as “the emotion most essential to the development of conscience and moral behavior” in psychology (Izard, 1991) and as the “internalized voice of moral authority” in philosophy (Griswold, 2007). Human beings feel guilty for transgressions they are causally responsible for, i.e., when they realize that they are responsible for an action or omission that violates certain moral norms or that they fail to live up to others’ expectations (i.e., personal guilt) (Baumeister, Stillwell, and Heatherton,1994; Chang and Smith, 2011; Izard, 1991). Moreover, guilt can also be encountered in inter-group interactions (Vollberg and Cikara, 2018): individuals may feel guilty for transgressions committed by members of social groups they identify as in-group, even when he/she is not directly responsible for these transgressions. However, the psychological and neural basis of group-based (collective) guilt and its relation to personal guilt are poorly understood.

As individuals rarely engage in social interactions without social identity or association (Mesquita, Boiger, and De Leersnyder, 2016; Tajfel and Turner, 1986), social emotions arising from such interactions are often tainted by group identity and inter-group appraisals (Mackie, Smith, and Ray, 2008; Smith and Mackie, 2015). Well-known cases of group-based or collective social emotions have been widely debated and reflected upon theoretically (Tollefsen, 2003; Smiley, 2017), and have received considerable empirical investigation in the past decades (Branscombe, Slugoski, and Kappen, 2004; Doosje, Branscombe, Spears, and Manstead, 1998; Ferguson and Branscombe, 2014; Wohl, Branscombe, and Klar, 2006). In this line of research, the most frequently used method for inducing group-based guilt is scenario-based imagination or recall of historical events involving intergroup conflict (Brown, González, Zagefka, Manzi, and Čehajić, 2008; Doosje et al., 1998; McGarty et al., 2005). These studies demonstrated that group-based guilt could facilitate inter-group reconciliation (Doosje, Branscombe, Spears, and Manstead, 2004; Lickel, Steele, and Schmader, 2011), reduce prejudice towards out-group (Amodio, Devine, Harmon-Jones, 2007; Powell, Branscombe, and Schmitt, 2005), and the acceptance of in-group responsibility for transgressions is a critical antecedent of group-based guilt (Castano and Giner-Sorolla, 2006; Čehajić-Clancy, Effron, Halperin, Liberman, and Ross, 2011).

Although the scenarios-based approach has consistently shown that self-reported guilt is elicited and group-based responsibility is perceived when participants are reminded of in-group misdeeds (Brown et al., 2008; Doosje et al., 1998; McGarty et al., 2005), the psychological nature of this pattern remains elusive: (1) Does the self-reported guilt reflect genuine feelings or merely reflect what the participants find morally appropriate for them to express? That is, do the participants express guilt-like sentiments in the absence of genuine feelings of guilt, only to meet social expectations dictating the expression thereof? (2) How does the brain encode group-based guilt? Specifically, does group-based guilt share common neurocognitive processes with personal guilt?

According to Intergroup Emotion Theory (IET; Mackie et al., 2008; Smith and Mackie, 2015), which posits that group-based emotions are similar to individual-level emotions in most respects (Rydell et al., 2008), we should observe shared cognitive processes (e.g., responsibility acceptance) and brain responses underlying personal and group-based guilt. In contrast, according to Display Rules Hypothesis (Diefendorff and Richard, 2003; Matsumoto, 1993), expressing certain emotions are socially desirable or even morally required for individuals in specific contexts. Even when an individual does not genuinely experience the emotion, he/she will nevertheless display it in order to be socially accepted or avoid sanctions. The Display Rules Hypothesis predicts that the cognitive-emotional processes associated with personal guilt are disparate from those associated with group-based guilt. Thus, if this hypothesis is true, we should not observe similar brain activation patterns for personal guilt and group-based guilt. Instead, we would expect higher activations in brain structures related to social norms and self-control, such as lateral prefrontal cortex (LPFC) (Buckholtz, 2015; Crockett, Siegel, Kurth-Nelson, Dayan, and Dolan, 2017), for group-based guilt because people would force themselves to display an emotion they are not experiencing.

To test these hypotheses, we developed a paradigm that combines an interpersonal transgression task that induces guilt (Koban, Corradi-Dell’Acqua, and Vuilleumier, 2013; Yu, Hu, Hu, and Zhou, 2014) with a minimal group manipulation that induces group identity (Dunham, 2018; Otten, 2016). In this paradigm, we manipulated participants’ relationship with transgressors (i.e., in-group/out-group) and the objective responsibility of the participants for the transgression (i.e., commit (responsible)/observe (not responsible)) in a situation where harm is committed. Specifically, participants either observed two in-group (*In-group_ Observe*) or two out-group (*Out-group_ Observe*) members cause harm to an anonymous victim group, or they themselves together with either an in-group (*In-group_ Commit*) or an out-group (*Out-group_ Commit*) member directly caused harm to the victims. Then they rated their level of guilt (Experiment 1) or divided 20 *yuan* (∼3 USD) between themselves and the victim group (Experiment 2). The monetary allocation decision, which the participants believed was unknown to the victim group, was included as a measure of reparative motivation and has been shown to be a reliable readout of guilty feeling (Ketelaar and Au, 2003; Yu et al., 2014). We predicted that group-based guilt would manifest as participants feeling guiltier, allocating more money, and having stronger brain activations when in-group members, rather than out-group members, caused harm to the victims. In addition, because shared responsibility plays a crucial role in experiencing group-based guilt, we predicted that shared responsibility may mediate compensatory behaviors through group-based guilt. Additionally, by combining neuroscience measures with this novel paradigm that simultaneously elicits personal and group-based guilt, we adopted conjunction analysis to compare the neurocognitive processes underlying personal guilt and group-based guilt to tease apart the Intergroup Emotion Theory and the Display Rules Hypothesis (Experiment 2).

## Results

### Group-based guilt elicited by an interaction-based minimal group paradigm

In Experiment 1, six participants were recruited for each experiment session. In Experiment 2, one participant was scanned each time, who would meet five confederates upon arrival at the laboratory. Then they were randomly assigned to a yellow group and a blue group, each consisting of three members. After the group manipulation, the participants played multiple rounds of a dot-estimation game either with two in-group partners or two out-group partners (Fig. 1). They were informed that another group of participants (i.e., victims), whom they would not meet, might have to receive electric shocks depending on the performance of the participants and their two partners (Fig. 2). Post-task manipulation checks revealed that after the minimal group manipulation, the participants identified significantly more with their own group than the other group in all of our experiments (see *Supplementary Results for Experiments 1 and 2*), indicating successful in-group/out-group manipulation.

**Fig. 1.**
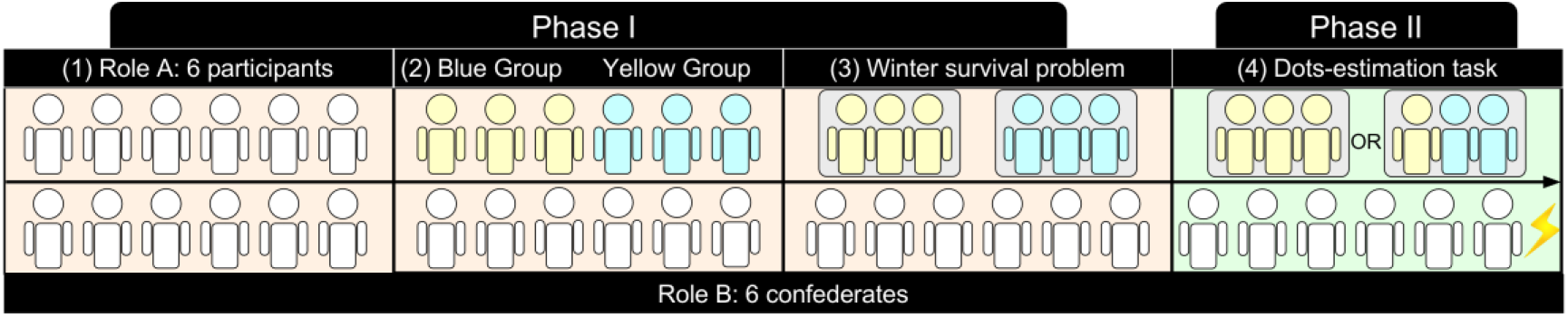
The timeline of the intergroup game. **(1)** Six same-sex students were recruited each time and were assigned Role A. They were told that another 6 participants (confederates) in another room were assigned Role B. **(2)** Then the 6 Role A participants were randomly divided into a ‘Yellow Group’ or ‘Blue Group’ to build the in-group/out-group context. They were also told that the participants in Role B were also divided into two 3-member groups. **(3)** To increase the feeling of affiliation to the new-built mini-group, each group solved a ‘winter survival problem’ within 6 minutes. **(4)** After mini-group manipulation, the participants (represented by the left-most figure of the three stick-figures) played a dot-estimation task with two in-group (e.g., the two yellow members in the example) or two outgroup partners (e.g., the two blue members in the example), and one group from Role B (the victims) would receive a painful shock if Role A (the transgressors) failed.

**Fig. 2.**
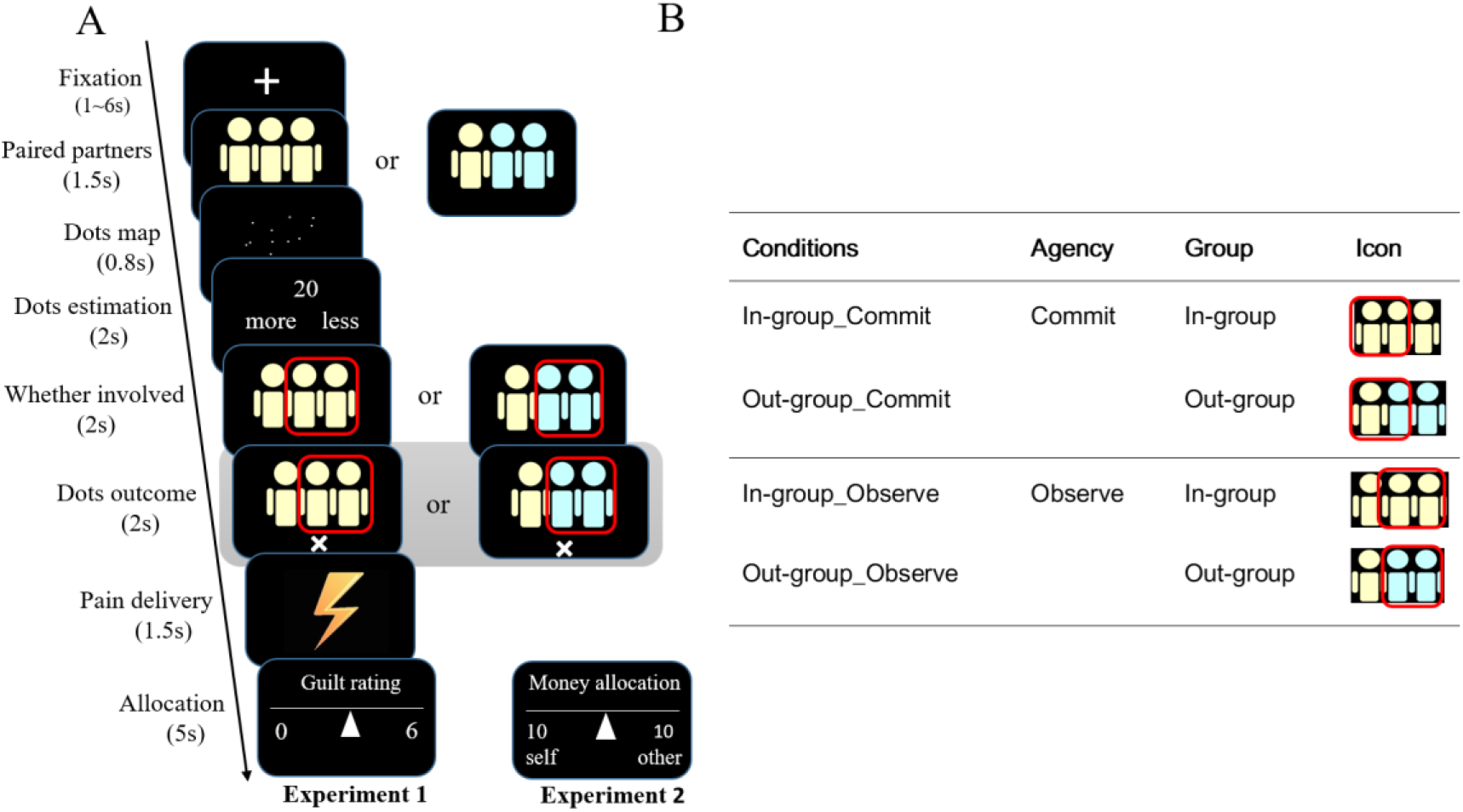
Experimental design and procedure. **(A)** Each trial began by informing the participant (the left individual) the in-group/outgroup partners he/she was paired with on the current round. Then a number of dots distributed randomly on the screen were presented. The participant needed to quickly estimate the number of the dots and compare the estimation with a specific number which appeared on the next screen. Then, two out of the three estimations were randomly chosen by two red rectangles and their correctness would be presented on next screen. If one or two of the chosen estimations were incorrect, a ‘×’ signal (indicating the failure of the current round) would be presented, and one group from Role B was randomly chosen to receive pain stimulation. After pain delivery, the participant rated feelings of guilt on a discrete 0-6 scale (Experiment 1) or allocated a portion of 20 *yuan* to victims (Experiment 2). The key screen for FMRI data analysis was the dot-outcome screen (red rectangle). **(B)** Different experimental conditions and the corresponding stick-figures presented in the procedure. In-group_ Observe: the estimations of the two in-group partners were selected, the estimation of the participant was not selected; Out-group_ Observe: the estimations of the two out-group partners were selected, and the estimation of the participant was not selected; In-group_ Commit: the estimation of one in-group partner was selected, as well as the estimation of the participant; Out-group_ Commit: the estimation of one out-group partner was selected, and also the estimation of the participant.

We used linear mixed effects models to examine how group membership (In-group *v.s.* Out-group) and objective responsibility (Commit *v.s.* Observe) modulated self-reported guilt (Experiment 1) and compensation behavior (Experiments 2). For both experiments, we found a significant interaction of group membership and objective responsibility on self-reported guilt (Experiment 1, Fig. 3A) and compensation behavior (i.e., monetary allocation; Experiment 2, Fig. 3B). Specifically, participants felt guiltier and allocated more to the victims in the *In-group_ Observe* condition than the *Out-group_ Observe* condition (Experiment 1: *t* = 5.10, *p <* 0.001; Experiment 2: *t* = 4.96, *p <* 0.001, see Table 1 and *Supplementary Results for Experiments 1and 2* for more details). This replicated previous findings from scenario-based studies on group-based guilt, namely, people report feeling more guilt for transgressions committed by members of their own group. The behavioral pattern was replicated in a direct replication of Experiment 2 (see *Supplementary Results for Experiments 3*).

**Fig. 3.**
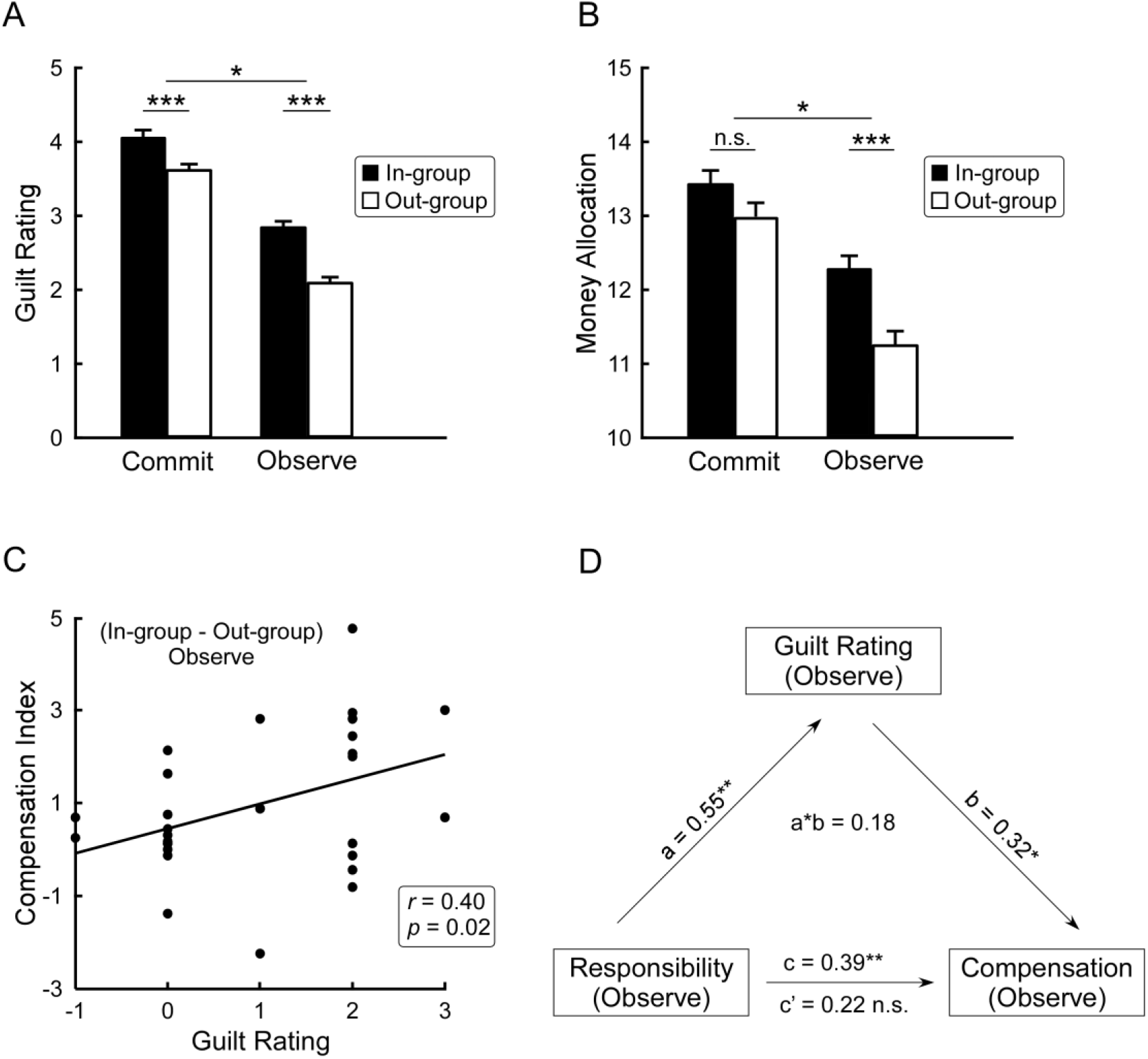
Behavioral results of Experiment 1 (A) and Experiment 2 (B). For Figures 2A and 2B, asterisks on the top of the graph indicate significant Group (In-group *v.s.* Out-group) by Objective responsibility (Commit *v.s.* Observe) interaction. Asterisks below indicate significance in post hoc test. **(C)** In Experiment 2, the post-experiment guilt rating difference (Observe: *In-group_ Observe > Out-group_ Observe*) positively correlated with the corresponding allocation difference. **(D)** The indirect pathway from the shared responsibility, via guilt rating, to compensation in Experiment 2. Error bars are standard errors estimated for data in each condition. *** *p* < .001, ** *p* < .01, * *p* < .05.

**Table 1.** Behavioral results in Experiments 1 and 2.

| Item | In-group_Commit | Out-group_Commit | In-group_Observe | Out-group_Observe | Interaction $T/F$ |
| --- | --- | --- | --- | --- | --- |
| **Experiment 1** |  |  |  |  |  |
| Online measure | | | | | Interaction $T$ |
| Guilt rating | 4.0 (.1) | 3.6 (.1) | 2.8 (.1) | 2.1 (.1) | 2.26* |
| Post-experiment measures | | | | | Interaction $F(1, 2)$ |
| Responsibility | 6.8 (.3) | 6.6 (.4) | 4.5 (.5) | 3.3 (.4) | 7.55* |
| Fear | 3.5 (.5) | 3.2 (.6) | 3.1 (.5) | 2.2 (.3) | 2.47 |
| Angry | 3.5 (.4) | 2.9 (.5) | 2.8 (.4) | 2.4 (.4) | 0.10 |
| **Experiment 2** |  |  |  |  |  |
| Online measure | | | | | Interaction $T$ |
| Monetary allocation | 13.5(.2) | 13(.2) | 12.3(.2) | 11.2(.2) | 2.14* |
| Post-experiment measures | | | | | Interaction $F(1, 3)$ |
| Responsibility | 6.9 (.3) | 4.4 (.4) | 6.6 (.3) | 3.5 (.4) | 5.41* |
| Guilt | 6.5(.3) | 5.9 (.4) | 4.1 (.4) | 3.2 (.4) | 1.05 |
| Fear | 3.6 (.3) | 2.7 (.4) | 3.3 (.3) | 2.6 (.3) | 0.16 |
| Angry | 3.1 (.4) | 2.7 (.4) | 3.0 (.3) | 2.6 (.3) | 0.11 |

*Note.* Standard errors (*SEs*) are shown in parentheses. *SEs* of Online measures of Experiment 1 and Experiment 2 were estimated by data from every single trial under each condition. *SEs* of Post-experiment measures were standard errors of means. Significant two-way interaction is denoted by \* $p < .05$ .

### Shared responsibility explains group-based guilt and compensation

To investigate the cognitive processes underlying group-based guilt, we examined the role of shared responsibility in group-based guilt. Not surprisingly, participants perceived higher responsibility in the *Commit* conditions than the *Observe* conditions, *F* (1, 23) = 151.17, *p* < 0.001, *η^2^_p_* = 0.87 for Experiment 1 and *F* (1, 30) = 65.01, *p* < 0.001, *η^2^_p_* = 0.68 for Experiment 2. Importantly, and supporting our hypothesis, this effect was modulated by partners’ group membership: the interaction between group membership (In-group *v.s.* Out-group) and objective responsibility (Commit *v.s.* Observe) was significant for both experiments (Experiment 1, *F* (1, 23) = 7.55, *p* = 0.011, *η^2^_p_* = 0.25; Experiment 2, *F* (1, 30) = 5.41, *p* = 0.03, *η^2^_p_* = 0.15; see Table 1 for details). Specifically, pairwise comparisons showed that the participants felt more responsible in the *In-group_ Observe* condition than in the *Out-group_ Observe* condition, *F* (1, 23) = 11.38, *p* = .003, *η^2^_p_* = 0.33 for Experiment 1 and *F* (1, 30) = 11.47, *p* < 0.001, *η^2^_p_* = 0.28 for Experiment 2. That they had no objective responsibility in either case suggests that the participants ‘inherited’ moral responsibility for transgressions committed by in-group members. Moreover, the difference in perceived responsibility between observing in-group versus out-group transgression was positively correlated both with the difference in self-reported guilt rating (Experiment1, *r* = 0.45, *p* = 0.03) and with the difference in monetary allocation (Experiment 2, *r* = 0.54, *p* = 0.002) between these two conditions. This indicated that the perceived moral responsibility was associated with the experience of group-based guilt. No significant effect was found for fear and angry emotion. The relationship between perceived moral responsibility in self-reported guilt was replicated in Experiment 3 (see *Supplementary Results for Experiments 3*).

Research on personal guilt has suggested that guilt functions as an intermediate state between acknowledging responsibility of transgression and reparative behavior (e.g., Yu et al., 2014). Here we provided more concrete evidence for this conjecture by a mediation analyses (Preacher and Hayes, 2008). We found a significant indirect path from perceived responsibility *via* self-reported guilt to monetary allocation (mediating effect estimate = 0.19, SE = 0.09, 95% confidence interval was [0.009, 0.357], see *Supplementary Results for Experiments 1 and 2* for detail). Importantly, for group-based guilt, we found a similar indirect pathway *via* guilt (the mediating effect estimate = 0.18, SE = 0.08, 95% confidence interval [0.003, 0.331] (Fig. 3D), see *Supplementary Results for Experiments 1 and 2* for detail). This finding lends support to the Intergroup Emotion Theory, according to which group-based guilt should function in a similar way as personal guilt in mediating the relationship between perceived responsibility and compensation behaviors.

### Brain activations associated with personal and group-based guilt

Here we presented the brain activation patterns revealed by the contrasts hypothesized to reflect group-based and personal guilt, respectively. The activation patterns corresponding to the main effects of Objective responsibility and Group can be found in *Supplementary Neuroimaging Results*.

To identify brain regions associated with group-based guilt, we focused on brain responses associated with the outcome feedback of dot estimation. We defined the critical contrast “*In-group_ Observe > Out-group_ Observe*”, which corresponds to the effect of group-based guilt. This contrast revealed activations in anterior middle cingulate cortex (aMCC; MNI coordinates = [6, 26, 28]; *k*= 85 voxels) and right anterior insula (AI; MNI coordinates = [42, 26, 4]; *k*= 78 voxels) (Fig. 4A). aMCC and right AI have been consistently implicated in imagining and experiencing personal guilt (Chang et al., 2011; Yu et al., 2014) and negative self-evaluation in social contexts (Immordino-Yang, McColl, Damasio, Damasio., 2009; Kédia et al., 2008; Koban et al., 2013; Sanfey, Rilling, Aronson, Nystrom, Cohen., 2003; Zaki, Ochsner, Hanelin, Wager, Mackey., 2007). To illustrate the activation patterns, we extracted the regional parameter estimates from 27 voxels around the peak coordinates at aMCC and right AI. The parameter estimates extracted from aMCC (Fig. 4B) and right AI (Fig. 4C) exhibited a pattern similar to the pattern of monetary allocation. Moreover, aMCC activation difference (*In-group_ Observe > Out-group_ Observe*) positively correlated with the post-scan guilt rating difference between these two conditions (*r* = 0.45, *p* = 0.011), indicating that the aMCC was involved in the processing of group-based guilt.

**Fig. 4.**
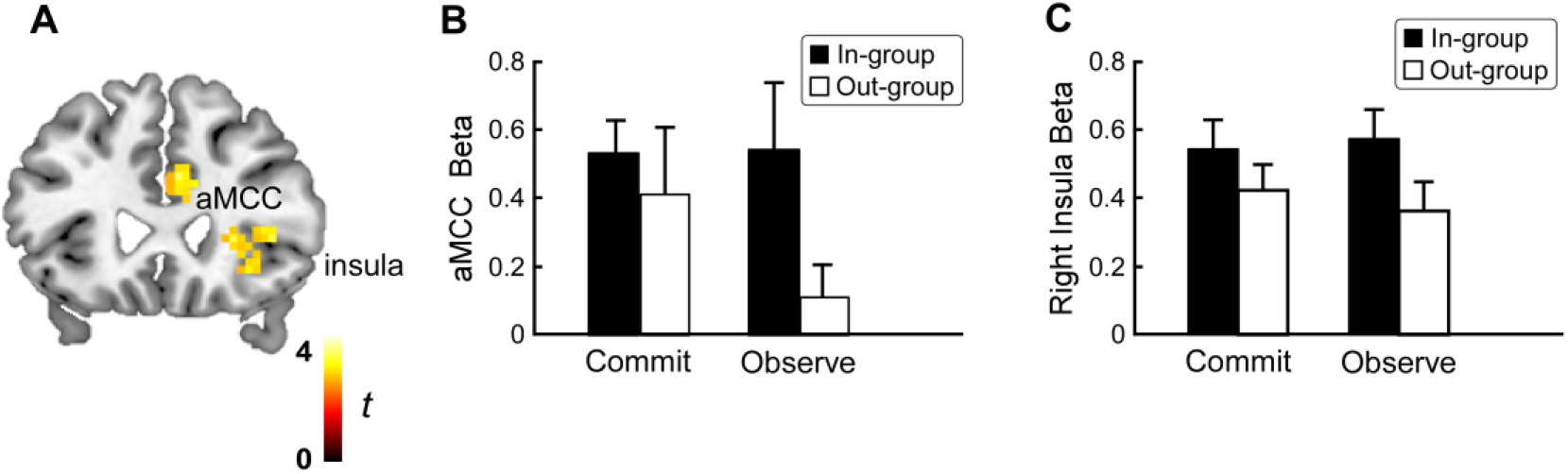
Brain activations related to group-based guilt. Results of the contrast ‘*In-group_ Observe > Out-group_ Observe*’ are shown in yellow-to-red clusters (A). Statistical parametric map was displayed at *P* < 0.005 uncorrected at peak level with cluster size ≥ 46 voxels. Regional activation patterns (i.e., beta estimates) were extracted from aMCC (B) and rAI (C) regions-of-interest (27 voxels around the peak coordinates of aMCC (MNI coordinates = [16, 26, 28]) and rAI (MNI coordinates = [33, 23, −2])). Error bars represent standard errors of the means.

We next examined whether group-based guilt shared a similar neurocognitive process with personal guilt. In the current study, ‘personal guilt’ was defined by the contrast ‘*Out-group_ Commit > Out-group_ Observe*’. This contrast, while keeping the impact of group membership to its minimal, captured the difference in the participants’ causal contribution to the transgression, thereby reflecting neural processing of personal guilt. Replicating previous neuroimaging findings about personal guilt (Koban et al., 2013; Yu et al., 2014), this contrast (*Out-group_ Commit > Out-group_ Observe*) revealed the activations in aMCC (MNI coordinates = [12, 17, 40]; *k*= 153 voxels) and supplementary motor area (SMA) (MNI coordinates = [15, 5, 67]; *k*= 62 voxels) (Fig. 5A).

**Fig. 5.**
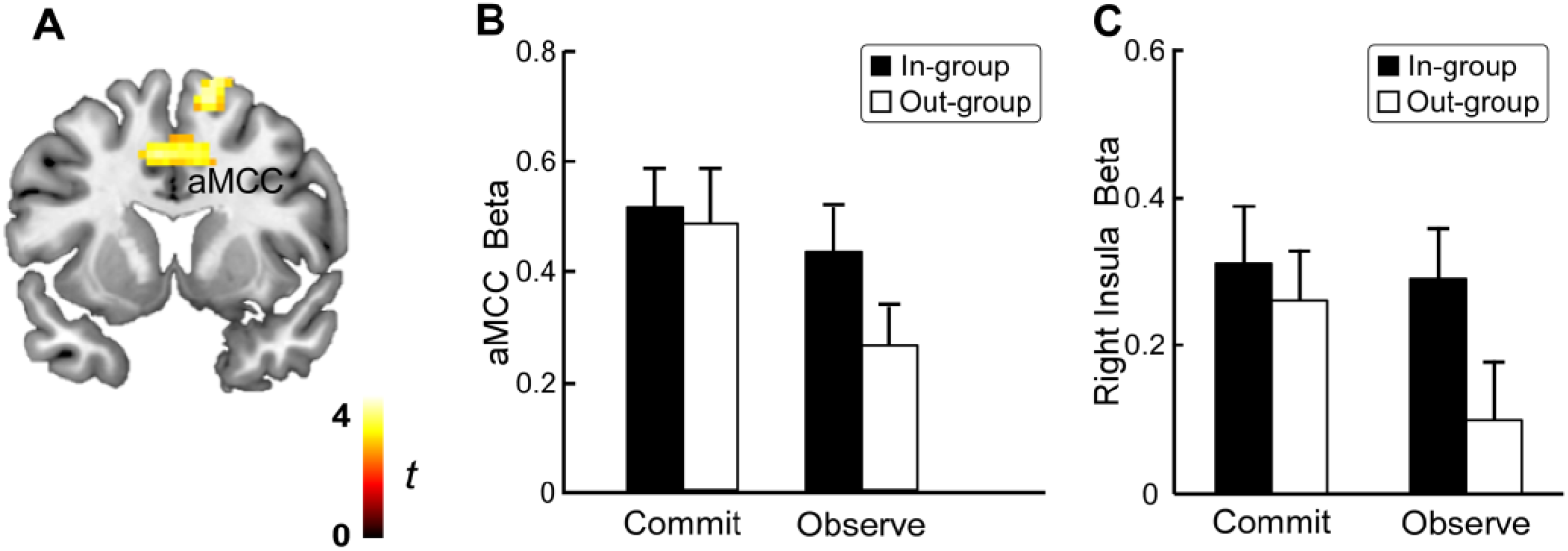
Brain activations related to personal guilt (‘*Out-group_ Commit > Out-group_ Observe*’). (A). Results of the contrast ‘*Out-group_ Commit > Out-group_ Observe*’ are shown in yellow-to-red clusters. Threshold for display was *P* < 0.005 uncorrected at peak level with cluster size ≥ 46 voxels. **(B).** Regional activation patterns (i.e., beta estimates) of aMCC was extracted from 27 voxels around the peak coordinates of aMCC (MNI coordinates = [12, 17, 40]). (C). Regional activation patterns of rAI was extracted from an independently defined region of interest (27 voxels around the peak coordinates of rAI reported in Yu et al., 2014; MNI coordinates = [36, 30, −8]). Error bars represent standard errors of the means.

We extracted the regional parameter estimates from 27 voxels around the peak coordinates of aMCC (Fig. 5B) and further demonstrated that the aMCC activation difference (*Out-group_ Commit > Out-group_ Observe*) was positively correlated with the difference in self-reported guilt between these two conditions (*r* = 0.45, *p* = 0.012). The right AI has been implicated in representing personal guilt (Kédia et al., 2008; Koban et al., 2013; Yu et al., 2014). We extracted the regional parameter estimates from 27 voxels around the peak coordinates of an indepdendetly defined right AI region of interest ([36, 30,-8], coordinates reported in Yu et al., 2014). The activations within this ROI (Fig. 5C) showed a similar pattern with that of the aMCC. Statistical analysis further confirmed that for this ROI, the activation in the *Out-group_ Commit* condition is stronger than in the *Out-group_ Observe* condition, *t* = 2.75, *p =* 0.01.

### Group-based guilt shares brain representation with personal guilt

The aMCC was implicated in the whole-brain contrasts both for the group-based guilt contrast (*In-group_ Observe* > *Out-group_ Observe*) and for the personal guilt contrast (*Out-group_ Commit > Out-group_ Observe*). This suggested the possibility of a shared neuropsychological processes underlying these two forms of guilt. To examine this in a more principled way, we first performed a conjunction analysis between the aforementioned contrasts. The overlapping region identified from the conjunction analysis confirmed that the overlapped aMCC ((MNI coordinates = [6, 26, 28]; *k*= 31 voxels), Fig. 6A) was sensitive to both personal guilt and group-based guilt. Moreover, we performed a multivariate pattern analysis (MVPA) as a supplementary analysis in the overlapped aMCC region to further test whether the neural representations of group-based guilt was similar to those of personal guilt. We demonstrated that the classifier trained on the personal guilt conditions within the overlapped aMCC region can distinguish different levels of personal guilt in a leave-one-subject-out cross-validation test (accuracy = 68% ± 6%, *p* < 0.001; Fig. 6B). More importantly, the classifier can also discriminate the two group-based guilt conditions (*Out-group_ Commit v.s. Out-group_ Observe*) with an accuracy of 71% ± 8%, *p* = 0.01 (Fig. 6B), demonstrating that the neural representations of personal guilt could discriminate those of group-based guilt.

**Fig. 6.**
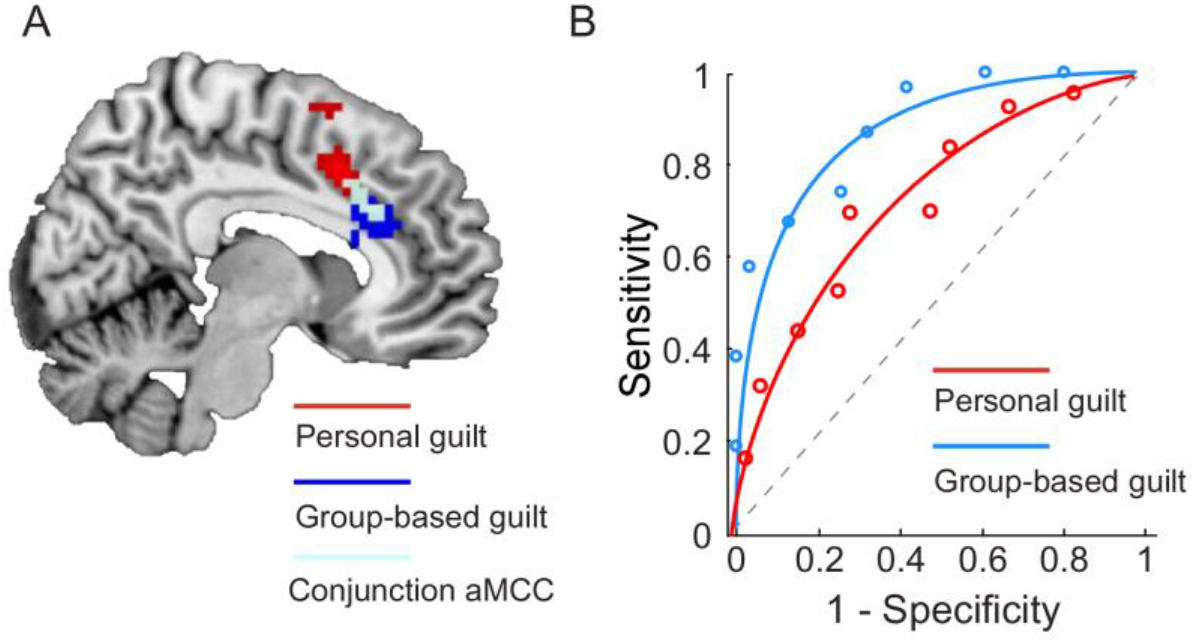
Results of multi-variate pattern analysis (MVPA). **(A)** Common areas for personal guilt and group-based guilt in aMCC. Threshold for display was *P* < .005 uncorrected at peak level with cluster size ≥ 30 voxels. **(B)** Receiver operating characteristic curves (ROC) for the two-choice forced-alternative accuracies. Red: Personal guilt; Blue: Group-based guilt.

## Discussion

In this study, we revealed neurocognitive profiles of group-based guilt and demonstrated its similarity with personal guilt. Specifically, shared responsibility for in-group transgressions is a crucial cognitive antecedent of group-based guilt just as personal, objective responsibility is crucial for personal guilt (Baumeister et al., 1994; Hoffman, 2001; Zahn-Waxler and Kochanska, 1990). Moreover, analysis of fMRI data suggested shared neuropsychological processes underlying these two forms of guilt. Our findings thus provide evidence for the Intergroup Emotion Theory account of group-based emotion (Mackie et al., 2008; Smith and Mackie, 2015), which posits that the neurocognitive machinery for individual-level emotions are co-opted in the group context.

Previous research on personal guilt has demonstrated that the sense of responsibility in causing suffering is closely related to the experience of guilt (Hoffman, 2001; Zahn-Waxler and Kochanska, 1990). Similarly, the perceived moral responsibility is also an important antecedent of group-based guilt (Čehajić-Clancy et al., 2011; Iyer et al., 2004). As demonstrated in the present study, although the participants had no personal contribution to transgressions in the *Observe* conditions, they shared greater moral responsibility for transgressions committed by in-group partners than out-group partners, and this vicarious responsibility was positively associated with the guilt rating (*In-group_ Observe > Out-group_ Observe*). These results are consistent with the shared responsibility account of group-based guilt (Smiley, 2017; Tollefsen, 2006): that individuals who identify themselves with a group acting badly would feel jointly responsible for the bad behaviors of the group. According to social identity theory (Tajfel and Turner, 1986), when people categorize themselves as a member of a group, and internalize the group component into self-concept, the actions of other in-group members will have a direct influence on self-perception. Thus, this ‘vicarious’ moral responsibility for in-group transgressions may result from the shared group membership with the harm-doers (Tajfel and Turner, 1986; Wohl et al., 2006). In a broader sense, these results suggested that responsibility, inasmuch as it is relevant for appraisals of social emotions, need be construed more broadly to include the sense of responsibility an individual inherited from their group identities. More importantly, our study, perhaps for the first time, demonstrates that similarities between group-based and personal guilt goes beyond self-reported emotional experience; the similarity is rooted in the neurobiology of emotional appraisal, lending support to the Intergroup Emotion Theory (Smith, 1993; Smith and Mackie, 2015). According to this theory, group members feel emotions in response to events affecting other in-group members as if those events were happening to themselves. An implication of this theory is that group-based emotions and personal emotions may share common neurocognitive substrates (Rydell et al., 2008). Our findings provide direct neural evidence for this hypothesis: aMCC responses to an in-group partners’ transgressions (relative to an out-group’s transgression) was positively associated with increased ratings of guilt, suggesting that aMCC is involved in experiencing group-based guilt.

Our findings shed new light on the role of aMCC in social-affective processing, extending its functional significance to inter-group processes (Lamm, Decety, and Singer, 2011; Shackman, Salomons, Slagter, Fox, Winter and Davidson, 2011). Thus, the aMCC activations here for transgressions made by an in-group partner may reflect the vicarious ‘personal guilt’ elicited by the perceived moral responsibility for transgressions, and raising the possibility of a shared neuropsychological process underlying these two forms of guilt (Woo et al., 2014). This possibility was further corroborated by our multivariate analyses (Chang et al., 2015; Wager et al., 2013) examining the similarity of neural representations of personal guilt and group-based guilt: our classifier trained on the personal guilt conditions could dissociate the group-based guilt conditions, suggesting that the group-based guilt could be developed on the basis of personal guilt.

Scenario-based imagination or recall of historical events involving intergroup conflict have been widely used to induce group-based guilt in previous studies (Branscombe et al., 2004; Brown et al., 2008; Doosje et al., 1998; McGarty et al., 2005), which laid a foundation for further research on group-based guilt. However, this approach’s strength of high ecological validity often comes at the cost of a well-controlled experimental manipulation, for example, by confounding group-based guilt with the level of harm inflicted on the victims (because the critical comparison in those studies is typically between a scenario depicting inter-group harm and a scenario depicting no harm). Thus, emotion measures obtained in these studies may be moderated by different levels of empathy or compassion for the victims (Branscombe et al., 2004; Brown et al., 2008; Doosje et al., 1998; McGarty et al., 2005). To address these pitfalls, in the current study, we assessed group-based guilt by comparing transgressions committed by in-group partners with identical transgressions committed by out-group partners (*In-group_ Observe > Out-group_ Observe*). Because the nature and extent of transgression is matched, any differential effect can thus be attributed to group membership (i.e., in-group/out-group).

Besides, the historical scenario-based method used in previous research limits the investigation of group-based guilt to the populations with a history of intergroup conflict (e.g., the intergroup conflicts between Israeli and Palestinians), which hinders the generalizability of the empirical evidence for conceptualizing group-based guilt as a universal psychological capacity. A strength of the interaction-based minimal group paradigm is that it extends the investigation of group-based guilt into populations without historical intergroup conflict. More importantly, the use of a minimal group paradigm provides a flexible experimental tool that could induce the intergroup emotion in a live and dynamic laboratory setting, thereby enabling a well-controlled laboratory study on group-based guilt to be conducted in natural circumstances. The investigation of the neuroscience of intergroup emotion is a relatively recent development within social psychology, but one that reflects a rapidly expanding interest in the interface between emotions and intergroup relations (Vollberg and Cikara, 2018). Following this trend, our interaction-based mini-group paradigm provides a flexible experimental tool to investigate the neural basis of group-based guilt.

## Conclusion

By combining fMRI with an interactive collective game, we provide behavioral and neural evidence demonstrating that group-based (collective) guilt and personal guilt share similar neurocognitive mechanisms. These findings provide neural evidence for the hypotheses that members of a group feel emotions in response to events affecting other in-group members as if those events affected them personally (Smith, 1993; Smith and Mackie, 2015). We highlight the crucial role of perceived responsibility in group-based guilt, which may help design interventions aimed at reducing unnecessary and excessive guilt inherited from past generations or other group members. Our findings shed light on the brain processes through which group membership is integrated into emotion appraisal, bridging gaps between research on emotion and inter-group relationship.

## Materials and Methods

### Participants

For Experiment 1 (behavioral experiment), we recruited 24 participants (12 females, mean age 19.3 ± 0.8 years). For Experiment 2 (fMRI experiment), thirty-five right-handed participants completed the experiment, four of which were excluded due to excessive head motion (>3 mm), leaving 31 participants (19 females, mean age 21.3 ± 1.1 years) in data analysis. None of the participants reported any history of neurological or psychological disorders. Written informed consent was obtained from every participant before the experiments. This study was carried out in accordance with the Declaration of Helsinki and was approved by the Ethics Committee of the School of Psychological and Cognitive Science, Peking University.

### Experimental Design and statistical analysis

#### Procedures of Experiment 1 (behavioral)

This study uses a 2 Group membership (In-group *v.s.* Out-group) × 2 Objective responsibility (Commit *v.s.* Observe) within-participant design, with ‘Group membership’ refers to whether transgressors were from the same or different group with the participants and ‘Objective responsibility’ refers to whether participants directly involved in transgressions.

##### Overview

In the current experiment, six participants of the same sex were recruited on each experimental session (none of them had known one another before the session). Upon arrival, participants were told that all six of them were predetermined to be assigned to group A (Transgressor-group) and six further co-players (confederates of the experimenter) in another room were assigned to group B (Victim-group). The task consisted of two phases. In the first, minimal group manipulation phase, the six participants of group A were randomly divided into two sub-groups of three members each to build in-group/out-group context; in the second phase, the participants played multiple rounds of a dot-estimation game either with two in-group partners or two out-group partners. The victims would receive electric shocks depending on the performance of the participant and /or his/her partners (Fig. 1). The participants were explicitly told that the victims could not reciprocate the electric shocks.

##### Minimal group manipulation

In the first phase, the six participants of group A were randomly divided into two sub-groups of three members each (a “Yellow Group” and a “Blue Group”). They were asked to wear a yellow or a blue T-shirt corresponding to their group membership. Each sub-group was required to work together to solve a “winter survival problem”(Johnson and Johnson, 1991) to enhance group identity. The background of the winter survival problem is that the plane the participants took had crash-landed in the woods of a northeastern region in mid-January, and they were required to rank-order 10 items salvaged from the broken plane (a lighter, a chocolate bar, a gun, newspaper, etc.) according to their importance for survival. The 3 individuals needed to discuss the problem together and reach a single consensus ranking within 6 min. As a manipulation check, we asked the participants to complete a scale of psychological distance to their group (a modified version of Inclusion of Others; Aron, Aron, and Smollan, 1992) and a 6-item questionnaire of group identity (example items include “How much do you identify with the Yellow group?” and “To what extent do you feel strong ties with the Yellow group?”; Falk, Heine, and Takemura, 2014) immediately after the mini-group manipulation. Player As were explicitly told that player Bs were also divided into two 3-member groups.

##### Pain calibration

After the group manipulation, participants were told that the victims (i.e., the confederates) would receive pain stimulation if the participants failed in their dot-estimation task (see below). To familiarize participants with the pain stimulation, all the participants underwent a pain calibration procedure. An intra-epidermal needle electrode was attached to the left wrist of participants for cutaneous electrical stimulation (Inui et al., 2002). Pain calibration begun with 8 repeated pulses (with each pulse having 0.2 mA and lasted for 5 ms with a 10 ms inter-pulse interval). Then we gradually increased the intensity of each pulse until participants reported 7 on a scale from 1 (‘not painful’) to 8 (‘intolerable’). All participants reported that the pain stimulation rated as 7 was really painful. They were told that the victims underwent the same pain calibration procedure and that the victims would receive the pain stimulation they rated as 7 if the transgressors failed in the dot-estimation task.

##### Dot-estimation task

In this task, each round began by informing the participant (represented by the left of the three stick-figures in the screens with the figures in Fig. 2A) that the two partners he/she paired with in the current round (represented by the middle and right of the three stick-figures in the screens of Fig. 2A) were from the “Yellow Group” or the “Blue Group”. Each of the three players was required to estimate the number of dots presented on the screen, press a corresponding button to indicate whether their estimate was ‘More’ or ‘Less’ than a reference number presented on the screen (randomly chosen from 19, 20, 21, and 22), and then press a button to confirm their choice. If the participant failed to confirm his/her choice within 2 seconds, the current trial would start again with a different dots map presented. The participants were explicitly told that the average accuracy for the dot-estimation task is 75%, to make them believe that they could estimate correctly. Two out of the three responses in the current round were randomly selected, as indicated by two red rectangles, and the outcome (success or failure) was presented on the next screen (see below for details on the different combinations of response selections). If the chosen estimates were both correct (i.e., filler trials), an ‘O’ sign would appear on the screen indicating that the current trial was successful and no painful stimulation would be delivered. The current trial terminated there. If one or both of the selected estimations were incorrect (i.e., experimental trials, Fig. 2B), a ‘×’ sign indicating failure of the current round was presented and one victim group of group B would be randomly selected to receive pain stimulation, as indicated by a shock sign appearing on the screen. Then, the participant was asked to rate his/her level of guilt on a 0-6 scale (in increments of 1) by pressing a key to increase or decrease the rating before pressing the space bar to confirm his/her choice.

The experimental trials (i.e., failure trials) consisted of the four combinations of the two experimental factors, namely, Group membership (In-group *v.s.* Out-group) and Objective responsibility (Commit *v.s.* Observe), forming a 2× 2 within-participant design. ‘Group membership’ refers to the group membership of the two partners and ‘Objective responsibility’ refers to whether the participant’s performance was selected and therefore directly involved in causing harm to the victims. Hence there were four experimental conditions: 1) the estimations of the two in-group partners were selected, and the estimation of the participant was not selected (*In-group_ Observe*); 2) the estimations of the two out-group partners were selected, and the estimation of the participant was not selected (*Out-group_ Observe*); 3) the estimation of one in-group partner was selected as well as the estimation of the participant (*In-group_ Commit*); and 4) the estimation of one out-group partner was selected as well as the estimation of the participant (*Out-group_ Commit*). Similarly, there were 4 possible combinations for success trials corresponding to the four experimental conditions and they were the filler trials. Unknown to the participants, the outcome of each trial was predetermined by a computer program, ensuring that all the conditions had the same number of trials.

The experiment consisted of 84 trials (16 for each experimental condition, and 5 for each filler combination). Trials were presented in a pseudorandom order to each participant with the constraint that no more than three consecutive trials were from the same condition. After the experiment, each participant rated on a 9-point Likert scale (1 = ‘not at all’, 9 = ‘very strong’), indicating his/her responsibility, fear, and anger in the four experimental conditions. After the experiment, each participant received 50 yuan (∼7.7 USD) for participation.

#### Procedures of Experiment 2 (fMRI)

Procedures of Experiment 2 were identical to those of Experiment 1, except that 1) one participant was recruited each time for scanning, and the participant met five confederates (2 male, 3 female, 23.6 ± 1.3 years) upon arrival at the laboratory; 2) at the end of each estimation-failure round, instead of rating feelings of guilt, the participant was asked to divide 20 *yuan* (∼3 USD) between him/herself and the 3 player Bs who received the pain stimulation, with the knowledge that the player Bs were unaware of the existence of this money distribution. The monetary allocation decision was included as a measure of reparative motivation and has been shown to be a reliable readout of guilty feeling (Ketelaar and Au, 2003; Yu et al., 2014). Thus, the amount allocated to the player Bs was interpreted as a measure of compensation for electric shocks. After scanning, participants rated their responsibility, fear, anger, and guilt on a 9-point Likert scale (1 = not at all, 9 = very strong) for each of the four experimental conditions. At end of Experiment 2, one trial was randomly selected and actualized as an extra payment. Thus, the participants’ final payoff was the baseline payoff (100 *yuan,* approximately 15 USD) and the amount of allocation the participants left for themselves.

### Direct replication of the behavioral findings of Experiment 2

To confirm the stability of the behavioral patterns observed in the fMRI experiment, we performed a behavioral experiment (Experiment 3) with the same procedures as the fMRI experiment in an independent sample of 36 participants, but with 6 same-sex participants recruited each time as in Experiment 1. One participant was excluded due to a technical error, leaving 35 (23 female, mean age 21.7± 0.9 years) for data analysis.

### Behavioral data analysis

The behavioral data were analyzed using linear mixed effects models implemented in the R software environment with the Imer4 package (Baayen, Davidson, and Bates, 2008; Bates, Mächler, Bolker, and Walker, 2014). To check how the Group (In-group *v.s.* Out-group) and the Objective responsibility (Commit *v.s.* Observe) modulated the guilt rating (Experiment 1) or compensation behavior (Experiments 2 and 3), the current LMM included two fixed-effect variables (Group and Objective responsibility) and their possible interactions, and one random factor (participants).To control for any potential confounding effects, we added all fixed factors (Group, Objective responsibility, and Group × Objective responsibility) into random slopes to better generalize the LMM analysis (Barr et al., 2013).

For all the mediation analyses in our study, we bootstrapped the mediating effect 20,000 times using the SPSS version of the INDIRECT macro (http://www.afhayes.com/) developed by Preacher and Hayes (Preacher and Hayes, 2008) and obtained the bias-corrected 95% confidence interval of the indirect effects.

### Imaging Data acquisition

Imaging data were acquired using a Siemens 3.0 Tesla Prisma scanner at the Beijing MRI Centre for Brain Research at Peking University (China). T2*-weighted echo-planar images (EPI) with blood oxygenation level-dependent (BOLD) contrast were collected in 33 axial slices parallel to the anterior-posterior commissure line to ensure coverage of the whole cerebrum (matrix 64 × 64, in planar resolution). Images were acquired in an interleaved order with no inter-slice gap (TR = 2000 ms, TE = 30 ms, voxel size = 3.5 mm × 3.5 mm ×3.5 mm, field of view = 224 mm ×224 mm, flip angle = 90°). A high-resolution, whole-brain structural scan (1×1×1 mm^3^ isotropic voxel MPRAGE) was acquired before functional imaging.

### Imaging Data preprocessing

The fMRI images were preprocessed using Statistical Parametric Mapping software SPM8 (Wellcome Trust Center for Neuroimaging, London, UK). The first five volumes of each run were discarded to allow for stabilization of magnetization. The remaining images were slice-time corrected, motion-corrected, re-sampled to 3×3×3 mm^3^ isotropic voxels, normalized to the Montreal Neurological Institute (MNI) space, spatially smoothed using an 8-mm full width at half maximum Gaussian filter, and temporally filtered using a high-pass filter with 1/128 Hz cutoff frequency.

### Whole-brain General Linear Model Analyses

Whole-brain exploratory analysis based on the general linear model was conducted first at the participant level, and then at the group level. To examine the neural responses to transgression outcomes, the data analysis focused on the brain responses associated with the presentation of the dot estimation outcomes. At the participant-level statistical analysis, failed dots-estimation outcomes corresponding to the four experimental conditions and 18 other regressors were separately modeled in a General Linear Model (GLM). Separate regressors in GLM were specified for fMRI responses to:

- R1: Combined regressor of no interest (duration = 4.3 s), which consisted of the presentation of the in-group/outgroup partners whom the participant paired with (“paired partners” screen), the dot map, and participants’ dot-estimation responses (“dot estimations” screens);
- R2-R5: Cue for involvement screen (duration = 2 s), which was indicated by a red rectangle around 2 of the 3 silhouettes. Each of the four experimental conditions was modeled in a separate regressor;
- R6-R9: Failed dot-estimation outcomes (2 s), separately modeled for each of the four experimental conditions;
- R10-R13: Successful dot-estimation outcomes (2 s), separately modeled for each of the four filler conditions;
- R14: The presentation of pain shock cue (1.5 s);
- R15: Allocation screen, which required the participant to allocate 20 yuan between him/herself and the victims (5 s);
- R16: Missed trials (4.3 s);
- R17-R22: Six head motion parameters, which were modeled separately in the GLM to account for artifacts in the scanner signal.

All regressors were convolved with a canonical hemodynamic response function. At the group-level, the four beta maps corresponding to the four experimental conditions of each transgressor were fed into a flexible factorial design matrix.

At the group-level, we defined the following contrasts:

- group-based guilt: *In-group_ Observe* > *Out-group_ Observe*;
- personal guilt: *Out-group_ Commit > Out-group_ Observe*;
- main effect of objective responsibility: (*In-group_ Commit + Out-group_ Commit*) > (*In-group_ Observe + Out-group_ Observe*);
- main effect of group membership: (*In-group_ Commit + In-group_ Observe*) > (*Out-group_ Commit + Out-group_ Observe*);

We focused on the group-based guilt contrast (*In-group_ Observe* > *Out-group_ Observe*) and the personal guilt contrast (*Out-group_ Commit > Out-group_ Observe*) as they were the main concern of our study. We took the simple effect contrast ‘*In-group_ Observe > Out-group_ Observe*’ as the defining contrast for group-based guilt because the victim’s harm as well as participants’ causal contribution to the harm were identical in these two conditions; the only difference was participants’ relationship with the transgressor. In the current study, ‘personal guilt’ was defined by the contrast ‘*Out-group_ Commit > Out-group_ Observe*’. This contrast, while keeping the impact of group membership to its minimal, captured the difference in the participants’ causal contribution to the transgression thereby reflected neural processing of personal guilt. Conjunction analyses

To identify brain areas that are shared by personal guilt and group-based guilt, we performed a conjunction analysis (Price and Friston, 1997) over the personal guilt contrast (*Out-group_ Commit > Out-group_ Observe*) and the group-based guilt contrast (*In-group_ Observe* > *Out-group_ Observe*). Conjunction analysis allowed us to combine these two comparisons to look for areas shared by personal guilt and group-based guilt while simultaneously eliminating areas activated in only one of the comparisons. Thus, this approach both increases statistical power (relative to only looking at, for example, the personal guilt contrast or the group-based guilt contrast), while also eliminating comparison specific activations which may reflect idiosyncratic influences of one of these two contrasts (Price and Friston, 1997). This conjunction was formulated as conjunction null hypothesis (Friston et al., 2005; Nichols et al., 2005) and should therefore only yield activations that are significantly present in both original contrasts of the conjunction. The null hypothesis for “conjunction null hypothesis” is that “not all contrasts activated this voxel.” If the conjunction results are significant, the null hypothesis is rejected and the conclusion is that “all contrasts activated this voxel.” That is, conjunctions represent a logical ‘and’, requiring both contrasts to be separately significant for the conjunction to be significant.

### Multivariate pattern analysis of imaging data

Compared to the univariate analysis, the Multivariate pattern analysis (MVPA) could increase the amount of information that can be decoded from brain activity (i.e. spatial pattern). Thus, a supplementary MVPA analysis was carried out in the conjunction region to check whether the spatial pattern of group-based guilt similar to those of personal guilt. We used linear Support Vector Machine (SVM) (Friedman et al., 2001; Wager et al., 2013) to train a multivariate pattern classifier on personal guilt trials and apply the classifier to discriminate group-based guilt. With a leave-one-out cross-validation method, we calculated the accuracy of the SVM classifiers using the forced-choice test (Chang et al., 2015; Wager et al., 2013; Woo et al., 2014). We also calculated the accuracy for *In-group_ Observe v.s. Out-group_ Observe*. To be noted that, in case the SVM effects were driven solely by the effects of response amplitude already observed in the GLM analysis, the mean univariate response magnitude in the overlapped region was subtracted (Coutanche, 2013; Smith, Kosillo, and Williams, 2011)

## Acknowledgements

### Funding

This work was supported by National Natural Science Foundation of China (31630034) and by National Basic Research Program of China (973 Program: 2015CB856400). Hongbo Yu was supported by a Newton International Fellowship from the British Academy (NF160700).

### Author contributions

ZL, HY, YZ, XZ design the study; ZL, HY collected data; ZL, HY analyzed data; ZL, HY, TK, XZ wrote the paper.

### Competing interests

The authors declare that they have no competing interests.

### Data and materials availability

All data needed to evaluate the conclusions in the paper are present in the paper and/or the Supplementary Materials. Original materials can be obtained from the corresponding author.

## Supplementary Materials

### Experimental Design

To explore the neurocognitive mechanism of group-based guilt and its relationship with personal guilt, we developed a minimal group-based interpersonal game to induce and simultaneously measure group-based guilt and personal guilt. In this paradigm, the participants are either involved in harm to the victims, or merely observe in-group or out-group members cause harm to the victims (i.e., group-based guilt), forming a 2 Group (In-group *v.s.* Out-group) × 2 Objective responsibility (Commit *v.s.* Observe) within-participant design. For group-based guilt, by observing guilt rating and money allocation following harm caused by in-group/out-group members, we could trace how group membership shapes individuals’ emotion and social decision making. Moreover, by combining the game paradigm with functional MRI, we explore the neural substrates of group-based guilt, and examine whether there are common neurocognitive processes underlying group-based guilt and personal guilt.

### Supplementary Results for Experiments 1 and 2

#### Minimal group manipulation check

As a manipulation check, we first examined whether the participants felt closer to the in-group than to the out-group. In Experiment 1, paired sample t-test showed that the participants indeed felt closer to and had stronger identity with the in-group partners than the out-group partners: closeness, 4.7 ± 0.2 *v.s.* 2.7 ± 0.2, *t* (23) = 10.0, *p* < 0.001, *d* = 2.08; identity, 5.4 ± 0.2 *v.s.* 3.5 ± 0.2, *t* (23) = 9.8, *p* < 0.001, *d* = 1.99. This was also true for Experiment 2. The participants felt closer to and identified more with the in-group partners than the out-group partners: closeness, 4.4 ± 0.3 v.s. 2.7 ± 0.2, *t* (30) = 7.8, *p* < 0.001, *d* = 1.41; identity, 4.0 ± 0.2 v.s. 3.1 ± 0.2, *t* (30) = 4.3, *p* < 0.001, *d* = 0.77. These results confirmed the success of in-group/out-group manipulation.

#### Group-based guilt elicited by an interaction-based mini-group paradigm

We examined the patterns of online guilt rating (Experiment 1) and monetary allocation (Experiment 2) using linear mixed effects. We found that in Experiment 1, participants felt more guilty when harm was caused by in-group partners than by out-group partners, *β* = 0.29, *SE* = 0.06, *t* = 4.90, *p <* 0.001, and when they committed in the harm than when they just observed the harm, *β* = 0.68, *SE* = 0.08, *t* = 8.50, *p <* 0.001, (Table 1; Fig. 3A). The Group × Objective responsibility interaction (*β* = 0.08, *SE* = 0.04, *t* = 2.26, *p* = 0.03) showed that the participants felt more guilty in the *In-group_ Observe* condition than the *Out-group_ Observe* condition, *β* = 0.37, *SE* = 0.07, *t* = 5.10, *p <* 0.001; the participants also felt more guilty in the *In-group_ Commit* condition than the *Out-group_ Commit* condition, *β* = 0.21, *SE* = 0.06, *t* = 3.22, *p <* 0.001. The guilt difference in *Observe* conditions (*In-group_ Observe > Out-group_ Observe*, 0.74) was bigger than *Commit* conditions (*In-group_ Commit > Out-group_ Commit*, 0.42), demonstrating that the group membership exerted a larger modulation on guilt rating when the participants were not causally involved in harm than when they were.

The pattern of monetary allocation (compensation behavior) in Experiment 2 was similar to the pattern of guilt rating in Experiment 1(Fig. 3B). Participants compensated more when the harm was caused by in-group partners than out-group partners, *β* = 0.38, *SE* = 0.11, *t* = 3.53, *p <* 0.001, and when they themselves committed the harm than when they just observe the harm, *β* = 0.73, *SE* = 0.09, *t* = 7.44, *p <* 0.001. The Group × Objective responsibility interaction (*β* = 0.16, *SE* = 0.08, *t* = 2.14, *p* = 0.04) showed that the participants allocated more in the *In-group_ Observe* condition than in the *Out-group_ observe* condition, *β* = 0.63, *SE* = 0.13, *t* = 4.96, *p <* 0.001; the allocation difference between *In-group_ Commit* condition and *Out-group_ Commit* condition was not significant, *β* = 0.22, *SE* = 0.13, *t* = 1.71, *p >* 0.05. Moreover, the allocation difference in *Observe* conditions (*In-group_ Observe > Out-group_ Observe*) was positively associated with the corresponding post-scan guilt rating difference, *r = 0.40, p = 0.02* (Fig. 3C). The higher the guilt rating, the more allocation to the victims, suggesting that the monetary allocation was motivated by prosocial emotion (i.e., group-based guilt), further confirming that our online measure (i.e., monetary allocation) captured the affective responses elicited by the interactive task. All of the above patterns were replicated in the replication Experiment 3 (*see Supplementary Results of Experiment 3*).

#### Shared responsibility explains group-based guilt and reparation

Research on personal guilt and reparative behavior has provided hints that guilt functions as an intermediate state between acknowledging responsibility of transgression and reparative behavior (Koban et al., 2013; Yu et al., 2014). We conceptually replicated this pattern for personal guilt by a mediation analyses (Preacher and Hayes, 2008). For Experiments 2 and3, the correlation between the perceived responsibility and monetary allocation was not significant (*r* = 0.26, *p* = 0.11) in Experiment 2, and was significant (*r* = 0.37, *p* = 0.04) in Experiment 3. Thus, to confirm whether the perceived responsibility correlated with monetary allocation, we combined Experiment 2 and Experiment 3 together to increase sample size. The results showed that the perceived responsibility was positively correlated with monetary allocation *r* = 0.28, *p* = 0.009, and the guilt rating was both related to the perceived responsibility (*r* = 0.56, *p* < 0.001) and monetary allocation (*r* = 0.39, *p* = 0.001). We then conducted mediation analyses to examine whether the effects of the perceived responsibility on monetary allocation was mediated by guilt rating. Results revealed a significant mediation effect of guilt rating on the relationship between the perceived responsibility and monetary allocation: the mediating effect estimate = 0.19, SE = 0.09, 95% bias corrected confidence interval was [0.009, 0.357]. According to IET, group-based guilt should function in a similar way as personal guilt in mediating the relationship between perceived responsibility and reparation. The same mediation analyses was thus then performed for group-based guilt. Consistent with this hypothesis, after combining the Experiment 2 and Experiment 3 together, we found an intermediate role of guilt between acknowledging responsibility of transgression and reparative behavior: the mediating effect estimate = 0.18, SE = 0.08, 95% bias corrected confidence interval [0.003, 0.331], Fig. 3D.).

#### Supplementary Neuroimaging Results

The activations for the main effect of Objective responsibility [(*In-group_ Commit + Out-group_ Commit*) > (*In-group_ Observe + Out-group_ Observe*)] and the main effect of Group [(*In-group_ Commit + In-group_ Observe*) > (*Out-group_ Commit + Out-group_ Observe*)] are presented in Table S2.

### Supplementary Results of Experiment 3

#### Behavioral validation experiment (Experiment 3)

The Experiment 3 was conducted with exactly the same procedures and parameters as Experiment 2 to confirm the stability of the behavioral patterns observed in the fMRI experiment. The manipulation checks (i.e., the IOS scale and group identity questionnaire) showed that the participants felt closer to and had stronger identity with the in-group partners than the out-group partners: closeness, 4.7 ± 0.2 *v.s.* 2.8 ± 0.2, *t* (34) = 10.0, *p* < 0.001, *d* = 1.66; identity, 5.4 ± 0.2 *v.s.* 3.5 ± 0.1, *t* (34) = 14.9, *p* < 0.001, *d* = 1.67. These results were consistent with Experiments 1 and 2, suggesting that the creation of the in-group/out-group context was successful in Experiment 3.

To confirm the stability of the behavioral patterns observed in Experiment 2, we examined the online monetary allocation in Experiment 3, and found that its patterns was the same as the pattern in the fMRI experiment. Specifically, the participants compensated more when the harm was caused by in-group partners than out-group partners, *β* = 0.52, *SE* = 0.11, *t* = 4.68, *p <* 0.001 and when they themselves committed the harm than when they were not, *β* = 0.79, *SE* = 0.14, *t* = 5.63, *p <* 0.001 (see Fig. S1). The Group × Objective responsibility was significant, *β =* 0.11*, SE =* 0.05, *t =* 2.28, *p* = 0.02. Further tests showed that the participants allocated more in the *In-group_ Observe* condition than in the *Out-group_ Observe* condition, *β =* 0.63, *SE =* 0.13, *t =* 4.96, *p <* 0.001; the participants also allocated more in the *In-group_ Commit* condition than in the *Out-group_ Commit* condition, *β =* 0.41, *SE =* 0.11, *t =* 3.56, *p <* 0.001. The allocation difference in the *Observe* conditions (*In-group_ Observe > Out-group_ Observe*, 1.31) was bigger than the *Commit* conditions (*In-group_ Commit > Out-group_ Commit*, 0.84), suggesting that the group membership exerted a larger modulation on monetary allocation when in the *Observe* conditions than in the *Commit* conditions. Moreover, consistent with the fMRI experiment, the allocation difference in *Observe* conditions was positively associated with the corresponding post-scan guilt rating difference, *r =* 0.48, *p =* 0.003.

We also examined the role of the perceived responsibility sharing in experiencing group-based guilt in Experiment 3. Specifically, Consistent with Experiment 2, the Group membership by Objective responsibility two-way ANOVA analysis on the sense of responsibility was significant, *F* (1, 34) = 4.35, *P* = 0.04 (Table S1). Planned *t* tests showed that participants sensed a higher level of responsibility in the *In-group_ Observe* condition than in the *Out-group_ Observe* condition, *t* (34) = 28.75, *p* < 0.001, and the difference of the perceived shared responsibility between these two conditions was also positively correlated with the corresponding guilt rating difference, *r =* 0.41, *p =* 0.01. No significant effect was found for fear and angry emotion.

---

**Fig. S1.**
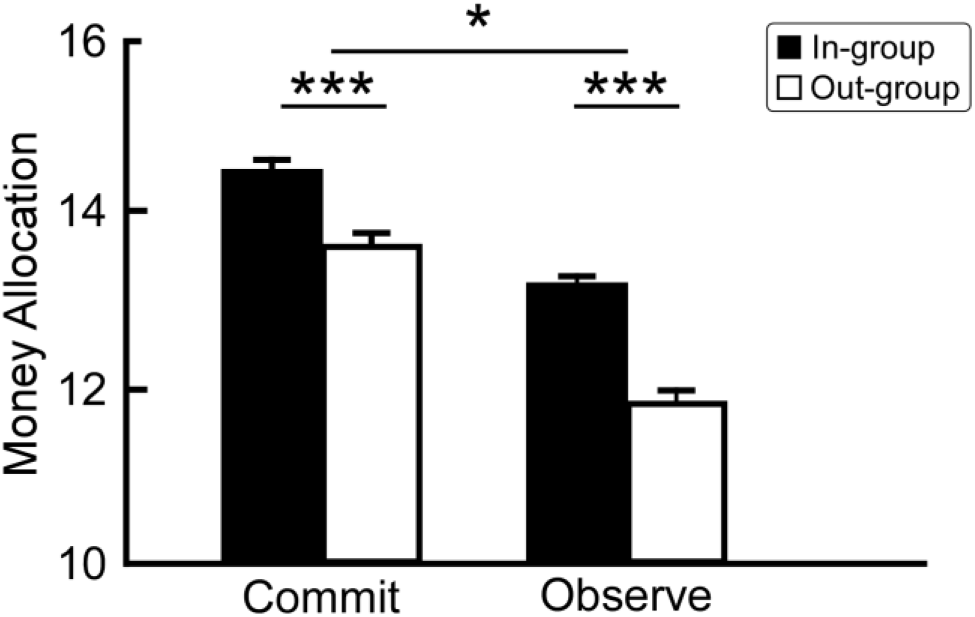
Behavioral results of Experiment 3. The monetary allocation in the four conditions in Experiment 3. For Figure S2-A, asterisks on the top of graph indicate significant Group (*In-group v.s. Out-group*) by Objective responsibility (*Commit v.s. Observe*) interaction. Asterisks below indicate significance in the post-hoc test. Error bars are standard errors estimated for data in each condition. *** *p* < .001, ** *p* < .01, * *p* < .05.

**Table S1.**
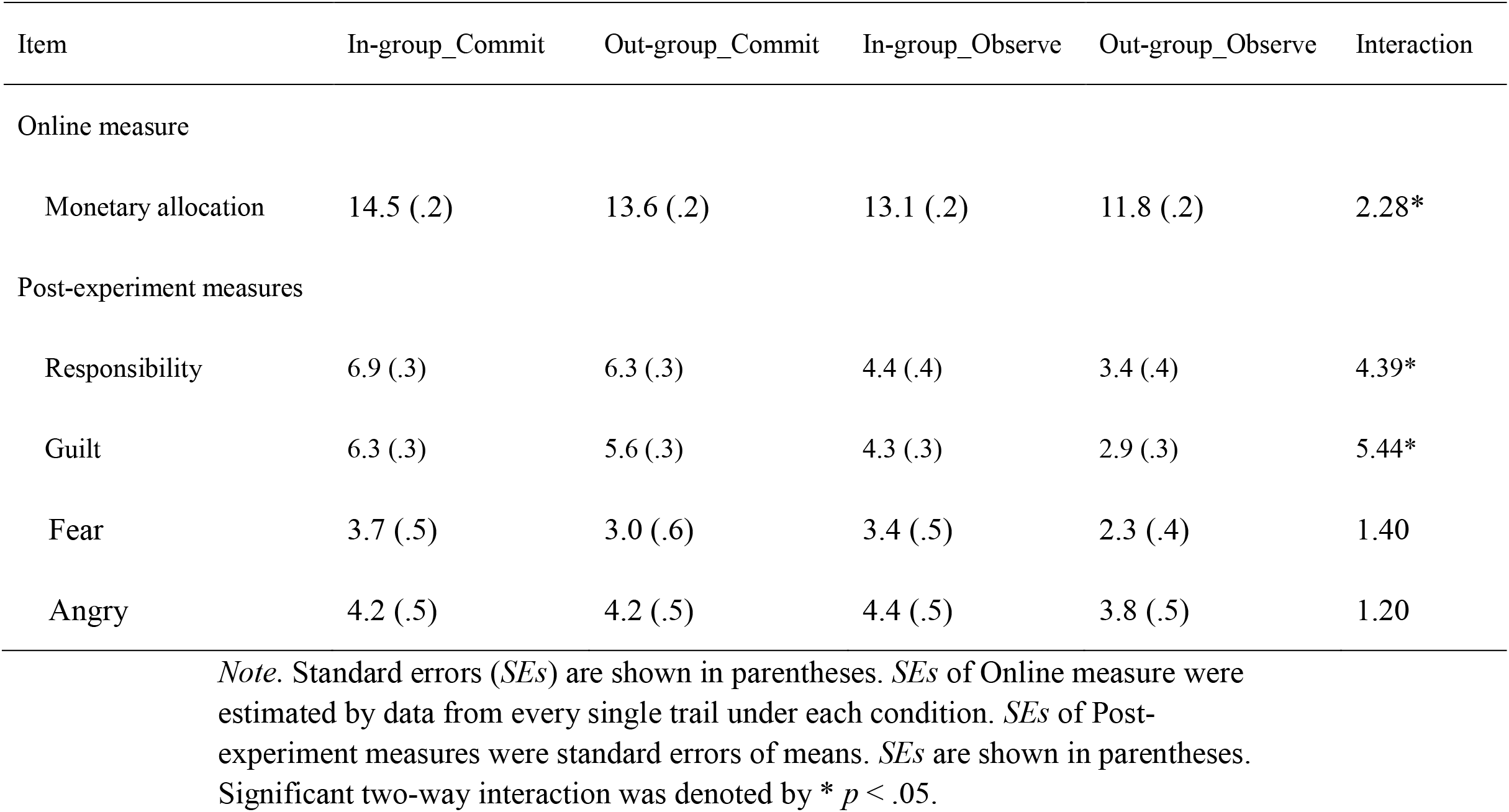
Results of Experiment 3.

| Item | In-group_Commit | Out-group_Commit | In-group_Observe | Out-group_Observe | Interaction |
| --- | --- | --- | --- | --- | --- |
| Online measure |  |  |  |  |  |
| Monetary allocation | 14.5 (.2) | 13.6 (.2) | 13.1 (.2) | 11.8 (.2) | 2.28* |
| Post-experiment measures |  |  |  |  |  |
| Responsibility | 6.9 (.3) | 6.3 (.3) | 4.4 (.4) | 3.4 (.4) | 4.39* |
| Guilt | 6.3 (.3) | 5.6 (.3) | 4.3 (.3) | 2.9 (.3) | 5.44* |
| Fear | 3.7 (.5) | 3.0 (.6) | 3.4 (.5) | 2.3 (.4) | 1.40 |
| Angry | 4.2 (.5) | 4.2 (.5) | 4.4 (.5) | 3.8 (.5) | 1.20 |

*Note.* Standard errors (*SEs*) are shown in parentheses. *SEs* of Online measure were estimated by data from every single trail under each condition. *SEs* of Post-experiment measures were standard errors of means. *SEs* are shown in parentheses. Significant two-way interaction was denoted by \* $p < .05$ .

**Table S2.** Brain activations revealed by univariate contrast (voxel-vise P < 0.005, minimum cluster extent = 46 voxels).

| Regions | Laterality | Peak MNI |  |  | Max | Cluster |
| --- | --- | --- | --- | --- | --- | --- |
|  |  | coordinates |  |  | T-value | size(voxels) |
|  |  | x | y | z |  |  |
| Group effect: In-group > Out-group |  |  |  |  |  |  |
| OFC | L | -27 | 47 | -11 | 4.3 | 161 |
| Left AI | L | -36 | 14 | -11 | 3.9 | 160 |
| aMCC | L,R | -6 | 26 | 34 | 3.6 | 200 |
| Right AI | R | 36 | 17 | -11 | 3.9 | 239 |
| Postcentral | R | 48 | -25 | 52 | 3.4 | 135 |
| SMA | R | 12 | 14 | 58 | 3.3 | 66 |
| Observe > Commit: |  |  |  |  |  |  |
| Cuneus | L,R | 0 | -61 | 25 | 5.3 | 590 |
| STS | L | -60 | -4 | -26 | 4.8 | 158 |
| TPJ | R | 54 | -61 | 22 | 4.1 | 332 |
|  | L | -45 | -70 | 28 | 4.3 | 226 |
| DLPFC | L | -24 | 23 | 46 | 4.7 | 274 |
| VMPFC | L,R | 0 | 56 | -17 | 3.9 | 226 |
| VLPFC | L | -57 | 29 | 4 | 3.8 | 163 |
| Commit > Observe: |  |  |  |  |  |  |
| aMCC | L,R | -3 | 8 | 43 | 4.3 | 476 |

*Note.* MNI coordinates are reported for peak activation.
R= right; L= left; STS = superior temporal sulcus; TPJ = temporoparietal junction; DLPFC = dorsolateral PFC; VMPFC = ventromedial PFC; VLPFC = ventrolateral PFC.

